# Genetic description of *Manania handi* and *Manania gwilliami*

**DOI:** 10.1101/747899

**Authors:** Mark A. Hanson, Hannah E. Westlake, Louise R. Page

**Author notes:** Corresponding authors: M.A. Hanson, L.R. Page.

## Abstract

Staurozoa is an intriguing lineage of cnidarians bearing both polypoid and medusoid characters in the adult body plan. Miranda et al. (2016) provided a massive descriptive effort of specimen collection, sequencing, and character evolution. We also recently described the neuromusculature of two staurozoan species: *Manania handi* and *Haliclystus “sanjuanensis.”* We found that our *M. handi* samples genetically matched *Manania gwilliami* samples used in Miranda et al. (2016). Taking advantage of newly-deposited *M. gwilliami* sequence data, we confirm the identity of our *M. handi* samples, and provide additional sequence data for *M. handi* and *H. sanjuanensis* for future staurozoan identification efforts.

## Introduction

Staurozoa (aka “stalked jellyfish”) is an ancient lineage of cnidarians with a medusa-like bell attached to a substrate by a polyp-like stalk. This body plan presents an intriguing intermediate between the ancestral anemone-like polyp and the derived medusae of other jellyfish. Whether this body plan is akin to an ancestral pre-medusa ancestor is unclear, as initial analyses of mitochondrial genome data conflicted with nuclear data, placing Staurozoa as either sister to Medusazoa (1, 2), sister to Cubozoa alone (3), or as a member of a grand Cubozoa, Scyphozoa, Staurozoa clade (4). Phylogenetic inference using combined mitochondrial and nuclear data also yields different results depending on phylogenetic analysis method (5). Given conflicting phylogenetic signals, the placement of Staurozoa within Cnidaria awaits additional sequence data for more robust comparisons.

As a distinct lineage of cnidarians bearing both polypoid and medusoid characters in the adult body plan, Staurozoa offers a unique opportunity to understand the evolution of other cnidarian groups. Miranda et al. (5) produced a monumental effort of classification, sequencing multiple genetic barcoding regions of diverse staurozoan species. Their phylogenetic inference provided long-needed clarity regarding hypothesized synapomorphic traits used for staurozoan taxonomic description.

We recently provided extensive characterization of the neuromusculature for two Staurozoan species: *Manania handi* and *Haliclystus “sanjuanensis”* (6). Notably, Miranda et al. (5) did not include a *M. handi* sample in their analyses, using only two *Manania* species: *M. uchidai* and *M. gwilliami. Manania* handi and *M. gwilliami* are both native to the west coast of North America (7), and genetic type specimens for *M. gwilliami* used for sequence comparisons in Miranda et al. (5) came from samples collected in California (hereafter *M. gwilliami* (CA)). Curious to support our identification of *M. handi* and *H. sanjuanensis*, we performed Sanger sequencing of mitochondrial 16s and COI loci, and also nuclear 18s (SSU) and ITS (internal transcribed spacer) regions for our specimens.

## Results and Discussion

Sequence data supported our identification of *H. sanjuanensis* as our specimen consistently clustered with other *H. sanjuanensis* samples in phylogenetic analyses for 16s, COI, 18s, and ITS sequences (Figure 1 and not shown). However, surprisingly we found that all four loci of our *M. handi* specimen were ≥99.7% identical to *M. gwilliami* (CA). Since Miranda et al. (5) and our study (6), additional COI data from *M. gwilliami* samples collected in 2018 in Bamfield BC Canada were deposited in GenBank; Bamfield is geographically near our sampling location in Victoria BC Canada. We used this opportunity to confirm these putative identifications of *M. handi* and *M. gwilliami*.

**Figure 1:**
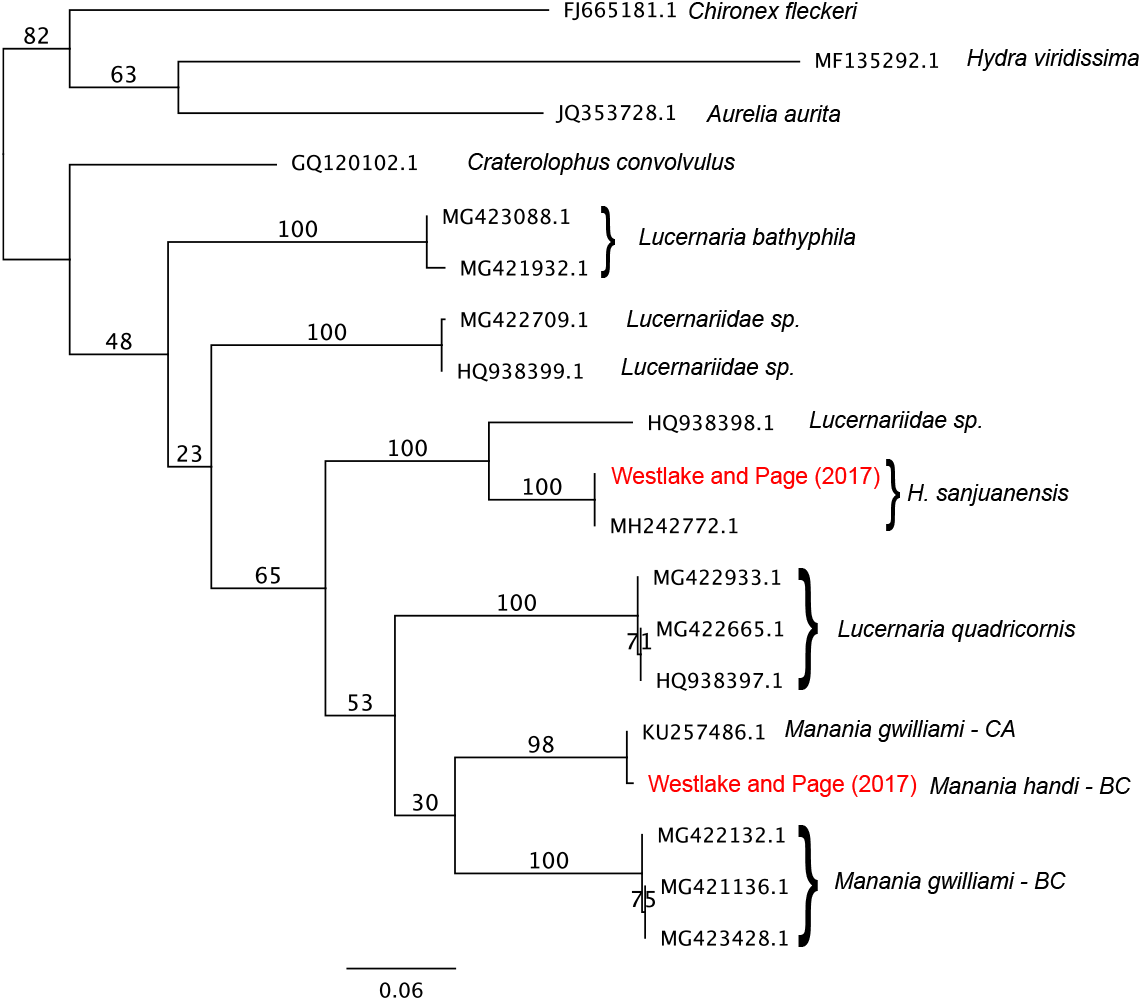
ML phylogeny (100 boostraps) of Staurozoan COI sequence, including samples from Westlake and Page (2017) in red. Notably, *M. handi* from Victoria (BC) and *M. gwilliami* (CA) form a distinct clade from *M. gwilliami* collected in Bamfield (BC). More robust phylogenetic inferences are discussed at length in Miranda et al. (5). GenBank accession numbers are given for pre-existing sequences.

We found that the three *M. gwilliami* samples from Bamfield clustered together as a separate group, distinct from our *M. handi* and the included *M. gwilliami* (CA) sample (Figure 1). While our specimen from Victoria BC is 99.7% identical at COI when compared to *M. gwilliami* (CA), our *M. handi* specimen was only 86.5% identical to *M. gwilliami* collected in Bamfield. We additionally confirmed our specimen as *M. handi* using characters described in Larson and Fautin (7) (Figure 2), and with Miranda et al. (personal communication). As such, we suggest that the type specimens for *M. gwilliami* used in Miranda et al. (5) were in fact *M. handi*. However, our analysis indeed suggests that *M. gwilliami* is a distinct species from *M. handi*, as *Manania* specimens collected in Victoria and Bamfield differ markedly at their COI locus. To further confirm *M. gwilliami* as a distinct genetic lineage, further sequence comparisons involving other *Manania* species should be performed.

**Figure 2:**
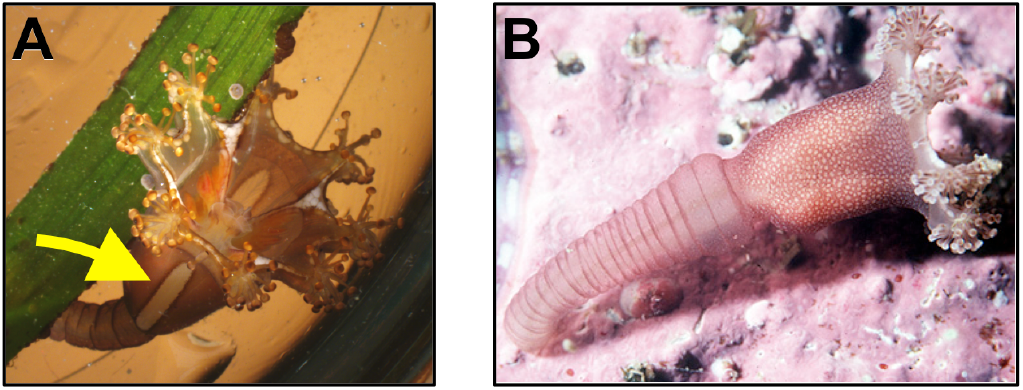
Representative images of *M. handi* (A) and *M. gwilliami* (B). Adult *M. handi* can be distinguished by distinctive “windows” of light-coloured pigment bounded by straight lines along the calyx (arrow), while adult *M. gwilliami* lacks these windows. The stalk of *M. gwilliami* is as long or longer than the calyx (7). *Manania gwilliami* photo by: Thomas Carefoot.

Species-level identifications are difficult for lesser-studied marine invertebrates, particularly uncultivable species like staurozoans. By performing an in-depth description in Westlake and Page (6) and generating supporting sequence data (this study), we provide morphological and genetic support for *M. handi* and *M. gwilliami* identification efforts.

## Materials and Methods

Samples were collected from the intertidal zone adjacent to Chinese Cemetery in Victoria BC, described fully in Westlake and Page (6). DNA was extracted from tissue samples using PrepMan Ultra. We used the 16s, 18s (SSU), COI and ITS primers from Miranda et al. (5). Sanger sequencing was performed by Macrogen USA. Sequences were aligned using MAFFT (8), and phylogenetic analyses were performed in Geneious R10 (9) using the PhyML plugin (10). Sequences generated in this study have been deposited in GenBank under the following accession numbers: MN329738-MN329745.

## Acknowledgements

HEW and LRP collected, photographed, and identified specimens. MAH sequenced specimens, analyzed data, and wrote this report. All authors read and approved the final report. Tom Carefoot generously provided the photograph of *Manania gwilliami* used in Figure 2. We owe special thanks to the team at the Centre for Biodiversity Genomics (University of Guelph, Canada) for making their *M. gwilliami* data publicly available.

